# Tracking animal reservoirs of pathogenic *Leptospira:* the right test for the right claim

**DOI:** 10.1101/2021.09.21.461181

**Authors:** Yann Gomard, Koussay Dellagi, Steve Michael Goodman, Patrick Mavingui, Pablo Tortosa

## Abstract

Leptospirosis, caused by a pathogenic *Leptospira* bacteria, is the most prevalent zoonosis worldwide and in this context has been extensively investigated through a One Health framework. Diagnosis of human leptospirosis includes molecular and serological tools, with serological Microscopic Agglutination Test (MAT) still being considered as a gold standard. Mammals considered as biological reservoirs include species or populations that are able to maintain chronic infection and shed the bacteria via their urine in the environment. *Leptospira* bacteria are often investigated using the same diagnosis tool, serological MAT. However, MAT testing of putative animal reservoirs can lead to mis-interpretations as it can signal previous infection and not necessarily bring in robust information regarding the capacity of such sero-positive animals to maintain chronic infection. We use previously published data and present new results on introduced and endemic small mammals to show that MAT should not be used for the identification of reservoirs. By contrast, serological data are informative on the level of exposure of animals occupying a specific environment. Finally, we present a sequential methodology to investigate human leptospirosis in a One Health framework that associates molecular detection in humans and animals, together with MAT of human samples using *Leptospira* isolates obtained from reservoir animals occurring in the same environment.

## Introduction

Leptospirosis is claimed as the most widespread bacterial zoonosis worldwide causing over one million human cases and nearly 60,000 deaths per year (Costa *et al*. 2015a). Despite its medical and veterinary importance, the burden of the disease remains underestimated in numerous countries, stimulating epidemiological investigations conducted in a One Health framework and aiming to identify the major drivers of the disease (Vinetz *et al*. 2005; Smythe and Chappel 2012; Allan *et al*. 2015). Leptospirosis is caused by pathogenic bacteria belonging to the genus *Leptospira* (family Leptospiraceae), which have been historically classified using antigenic determinants through Microscopic Agglutination Test (MAT) (Martin and Pettit 1918) and Co-Agglutination Absorption Technique (CAAT), allowing to define over 20 serogroups and 300 serovars (Levett 2001; Picardeau 2013). Molecular tools have been more recently developed and have revealed a high genetic diversity of *Leptospira* (Picardeau 2013; Saito *et al*. 2013; Bourhy *et al*. 2014) with several additional species later uncovered by genomic approaches (Guglielmini *et al*. 2019; Vincent *et al*. 2019).

The main biological cycle of pathogenic *Leptospira* involves wild or domestic animals acting as reservoirs through the shedding of the bacteria via their urine in the environment (Ko, Goarant and Picardeau 2009). Humans get mostly (but not only, see Bulach *et al*. 2006) infected through indirect contact with a contaminated environment. Although virtually all mammal species can get infected by these pathogenic bacteria, some requirements are needed to consider them as reservoirs (Babudieri 1958), and indeed only a limited number of species have been definitively shown to support chronic maintenance of the bacteria in their kidneys. Rodents are considered as the main reservoirs but other mammals such as bats (Dietrich *et al*. 2015), invasive (Cosson *et al*. 2014; Costa *et al*. 2015b) or endemic (Dietrich *et al*. 2014; Lagadec *et al*. 2016) small terrestrial mammals as well as cattle (Barragan *et al*. 2016; Guernier *et al*. 2016) have been identified as important reservoirs of *Leptospira*. The multiplication of pathogenic bacteria in animal reservoirs has been examined in experimental infections of mice under laboratory controlled conditions (Ratet *et al*. 2014). Using bioluminescent *Leptospira*, authors showed that a systemic infection associated with weight loss can occur within three days following intra peritoneal infection. Thereafter, within a week, bacteria become rapidly invisible while animals return to a body weight that is hardly distinguishable from that of control animals. Then, a bioluminescent signal of *Leptospira* appears in two spots, corresponding to kidneys where bacteria actively divide leading to a glowing signal persisting for months while systemic infection has apparently irreversibly vanished (Ratet *et al*. 2014).

Hence, the fate of pathogenic *Leptospira* appears to be different in reservoir and incident hosts, with a systemic infection followed by renal colonization in the former contrasting with a general absence of renal colonization in the latter. The separation between reservoir and incident hosts may be not that clear cut and depends on different parameters. Indeed, experimental infections have shown that survival of infected animals and shedding of bacteria depend on the bacterial strains, the infecting bacterial dose, the vertebrate species concerned, as well as the routes of infection (Ratet *et al*. 2014; Matsui *et al*. 2015; Wunder *et al*. 2016; Gomes-Solecki, Santecchia and Werts 2017). For instance, experimental infection of Golden hamsters considered as models of acute infection may lead to chronic shedding in the few animals surviving the infection (Cordonin *et al*. 2019). However, the colonization of renal tubules, which is typical of animal reservoirs, has a considerable immunological consequence: pathogenic *Leptospira* organized in biofilms in the lumen of renal tubules (Ristow *et al*. 2008) remain hidden from the immune system. Since the duration of sero-positivity following infection is not well known (Lloyd-Smith *et al*. 2007), the immunological signature detected in sera may not reflect the *Leptospira* that are chronically shed by the reservoir animal.

Beside its use in *Leptospira* classification, MAT is considered as the reference test for leptospirosis diagnosis in incident hosts (humans and domestic animals), as it allows detecting host antibodies testifying to current, recent, and past infections (Levett 2001; Musso and La Scola 2013). MAT has also been widely used for the investigation of animal reservoirs (Roberts *et al*. 2010; Desvars *et al*. 2012; Assenga *et al*. 2015; Andersen-Ranberg, Pipper and Jensen 2016; Rodrigues *et al*. 2016; Sigaud *et al*. 2009) but some studies indicate that MAT does not definitively verify the carrier status of a given animal species (Ellis, O’Brien and Cassells 1981; Miraglia *et al*. 2008; Libonati, Pinto and Lilenbaum 2017; Sant’anna *et al*. 2017). In the present work, we present further support for these latter observations and argue that the use of MAT may lead to misconclusions regarding the importance of investigated animal species as reservoirs.

To support our purposes, we focused on animal species known as pathogenic *Leptospira* reservoirs on Southwestern Indian Ocean (SWIO) islands. This region is home to a wide diversity of mammals, many being endemic, as well as introduced rodents (family Muridae) and shrews (family Soricidae). The typing of *Leptospira* excreted by mammals in this region has demonstrated high levels of *Leptospira*-host specificity (Dietrich *et al*. 2014, 2018; Gomard *et al*. 2016; Lagadec *et al*. 2016). Indeed, the region is home to a large diversity of bats from seven different families that represent multiple colonizations of the region and endemic terrestrial mammals of the family Tenrecidae and subfamily Nesomyinae (each representing separate adaptive radiations), which appear to be the exclusive reservoirs of specific pathogenic *Leptospira*, providing interesting biological circumstances to address the power of MAT for investigating leptospirosis epidemiology.

Three animal species from the SWIO region, known to host distinct bacterial lineages/species, were included in the present investigation. Using published molecular and serological data together with original results, we demonstrate that *Leptospira* serological signatures are not necessarily connected to the *Leptospira* excreted by animal reservoirs. We demonstrate the shortfalls of MAT used alone for the identification of *Leptospira* animal reservoirs and discuss the utility of MAT for clarifying leptospirosis epidemiology.

## Materials and methods

### Ethical considerations

Biological materials screened in the present study were sampled in the context of a research program for which permits numbers and IACUC acceptance have been presented elsewhere (Dietrich *et al*. 2018).

### Animal sampling, Leptospira serological, and molecular data

Three mammal species from the SWIO were included in the study: *Mormopterus acetabulosus* (Molossidae), an insectivorous bat species endemic to Mauritius Island; *Tenrec ecaudatus* (Tenrecidae), an omnivorous terrestrial mammal species endemic to Madagascar and introduced to several SWIO islands, including Reunion Island and Mayotte; an invasive rodent species, *R. rattus* (Muridae) sampled both on Reunion Island and on Mayotte (Table 1). Most of these species were previously investigated for *Leptospira* infection through molecular and/or serological methods by different researcher groups (Desvars *et al*. 2012, 2013; Pagès *et al*. 2015; Guernier *et al*. 2016; Lagadec *et al*. 2016; Dietrich *et al*. 2018). In addition, we produced serological data through MAT for the *M. acetabulosus* samples. The same individual specimens were previously investigated for *Leptospira* infection through molecular methods (Dietrich *et al*. 2018). MAT was performed essentially as previously described (Biscornet *et al*. 2017) using 18 *Leptospira* strains and screening most serogroups reported in both human cases and animals on SWIO islands (Table S1). A serum was considered as positive when the MAT titer was ≥ 1:100.

**Table 1.**
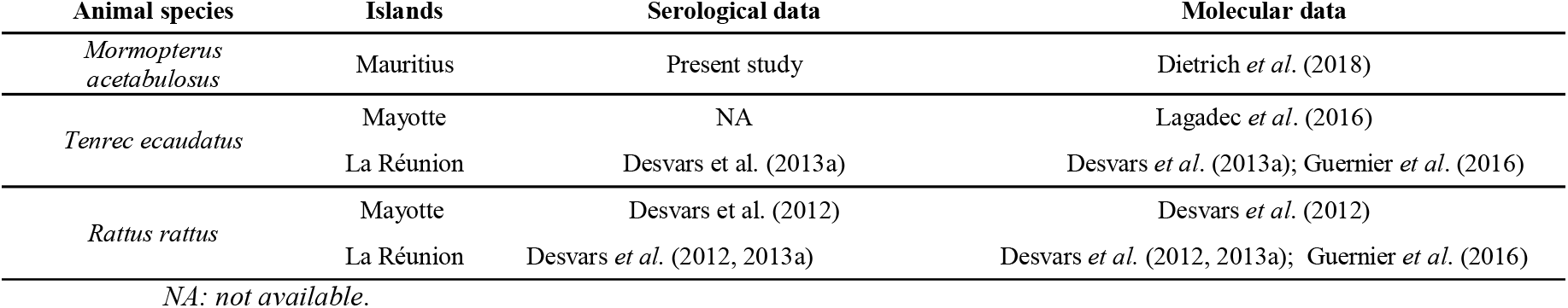
Animal species used in the present study and the associated publications for *Leptospira* investigations.

| Animal species | Islands | Serological data | Molecular data |
| --- | --- | --- | --- |
| *Mormopterus acetabulosus* | Mauritius | Present study | Dietrich *et al.* (2018) |
| *Tenrec ecaudatus* | Mayotte | NA | Lagadec *et al.* (2016) |
|  | La Réunion | Desvars *et al.* (2013a) | Desvars *et al.* (2013a); Guernier *et al.* (2016) |
| *Rattus rattus* | Mayotte | Desvars *et al.* (2012) | Desvars *et al.* (2012) |
|  | La Réunion | Desvars *et al.* (2012, 2013a) | Desvars *et al.* (2012, 2013a); Guernier *et al.* (2016) |

NA: not available.

## Results

### Bats

Serotyping of *Mormopterus acetabulosus* samples through MAT indicates that 20.0% (6/30) of specimens were seropositive, with sera agglutinating Panama and Pyrogenes serogroups (Table 2 and S2). Using nucleic acids extracted from the kidneys of the same individual specimens, Dietrich et al. (2018) reported that 73.3% (22/30) of the animals tested positive through Real-Time Polymerase Chain Reaction (RT-PCR), showing poor agreement between MAT and RT-PCR (Kappa test = 0.17). More specifically, the six MAT-positive bats were also positive by RT-PCR while, most importantly, 66.6% of the remaining MAT-negative bats (16/24) tested positive by RT-PCR. The sequencing of RT-PCR-positives specimens (also positive in MAT) confirmed that *M. acetabulosus* harbors a *Leptospira* bacterial sequence falling within the *Mormopterus*-borne *Leptospira* monophyletic clade embedded in *L. borgpetersenii* (Dietrich *et al*. 2018) (Table 2).

**Table 2.**
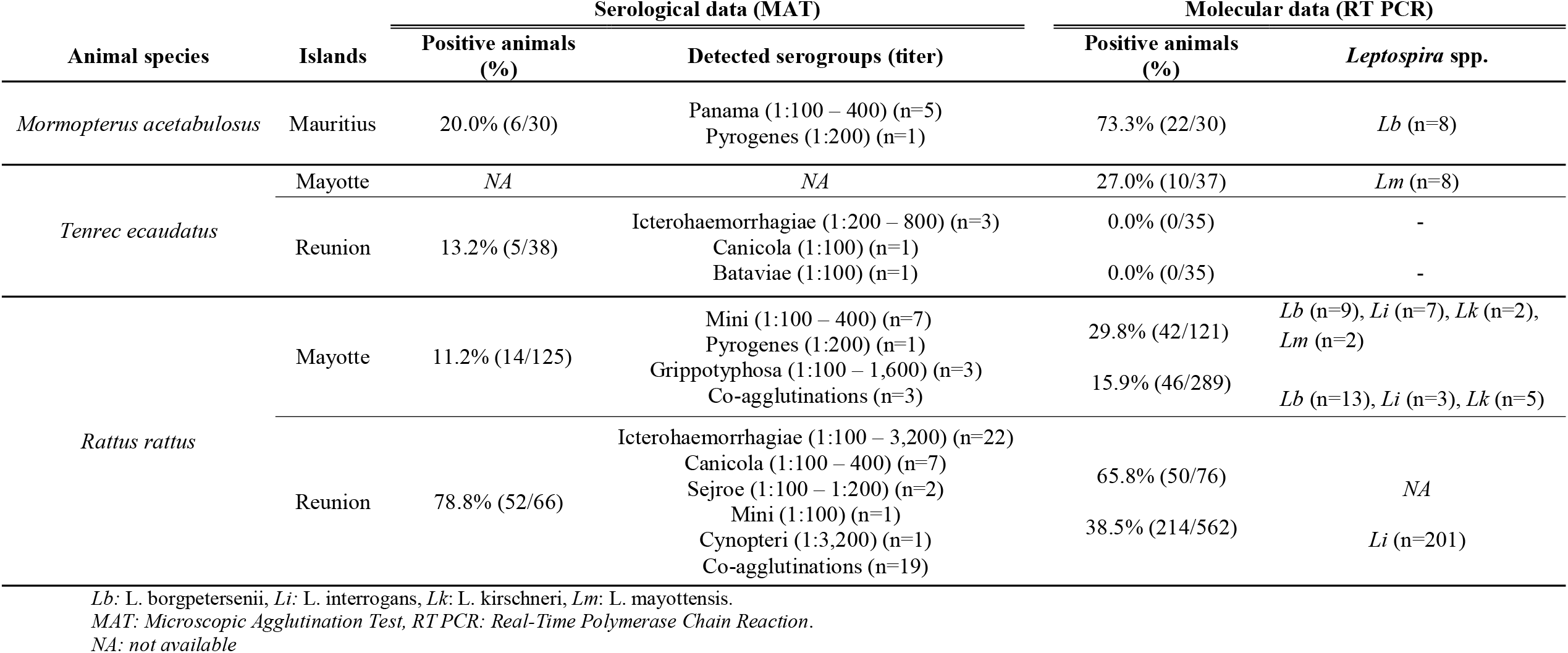
Comparison of serological and molecular *Leptospira* data obtained from the investigated animal species.

### Rats

Introduced populations of *Rattus rattus* are present on both Mayotte and Reunion Island, but molecular and serological screenings highlight striking differences between these two islands (Table 2). On Mayotte, three serogroups have been previously reported, namely Mini, Pyrogenes, and Grippotyphosa, whereas on Reunion Island the main detected serogroups correspond to Icterohaemorrhagiae, Canicola, Sejroe, Mini, and Cynopteri (Desvars *et al*. 2012, 2013). The molecular investigations of *R. rattus* on both islands confirms sharp inter island differences, with Reunion Island animals harboring strictly *L. interrogans* (Guernier *et al*. 2016), while on Mayotte this rodent may harbor either of three distinct *Leptospira* species (*L. interrogans, L. borgpetersenii*, and *L. kirschneri*) (Lagadec *et al*. 2016). On Reunion Island, a study investigated an outbreak of human leptospirosis after a triathlon and included 10 *R. rattus* that were incidentally trapped at the site where the sporting event took place a few weeks before the event. Five out of the 10 sampled rats tested positive by PCR based on kidney samples. The sequencing of the positive samples revealed only *L. interrogans* (Pagès *et al*. 2015; Guernier *et al*. 2016). Of note, two of the PCR-positive specimens were positive through MAT, whereas the remaining PCR-positive rats were sero-negative.

### Tenrecs

On Reunion Island, three serogroups have been reported in *Tenrec ecaudatus*: Icterohaemorrhagiae usually detected with high titers, while Canicola and Bataviae serogroups are agglutinated with low titers (Desvars *et al*. 2013a; Sigaud *et al*. 2009) (Table 2). Although these serogroups have been detected with moderate to high prevalence on Reunion Island, *T. ecaudatus* is not considered as a *Leptospira* reservoir on that island since renal carriage could not be demonstrated through two independent studies (Desvars *et al*. 2013; Guernier *et al*. 2016) (Table 2). This absence of infection on Reunion Island contrasts with the situation on Mayotte, where *T. ecaudatus* was identified as the exclusive reservoir of *L. mayottensis*, a pathogenic species commonly associated with human leptospirosis on that island (Lagadec *et al*. 2016) (Table 2).

## Discussion

Microscopic Agglutination Test (MAT) has been, and still is, considered the gold standard for leptospirosis diagnosis in humans. A meta-analysis has calculated the mean prevalence in reservoir mammals using the data published in 300 papers including eight different taxonomic orders (Andersen-Ranberg, Pipper and Jensen 2016). MAT and PCR were given an equivalent weight in that analysis, and the nature of the screened samples, *i*.*e*. blood (for MAT and PCR) or kidney/urine (for PCR only) was not taken into consideration. Hence, acute/passed infections and chronic kidney carriage were not distinguished in that study, as has been done in several others. In the bat samples screened in the present study, we demonstrate a poor agreement between data from serological and molecular analyses. Similar findings were also reported on a fruit bat species, *Pteropus alecto* (Pteropodidae), from Australia (Cox, Smythe and Leung 2005), which indicated poor agreement between PCR (detection on kidneys) and serological data; these results underlined that a carrier status for this species could not be shown based on serology. In Brazil, studies have reported limited or absence of agreement between PCR (detection in urine) and serological results in livestock animals or asymptomatic dogs (Hamond *et al*. 2014; Sant’anna *et al*. 2017).

The bat species investigated herein belongs to the genus *Mormopterus*, which includes within the SWIO region two other species, *M. francoismoutoui* and *M. jugularis*, endemic to Reunion Island and Madagascar, respectively. Recently, the research on these three molossid bats has shown that they shelter pathogenic *Leptospira* clustering into a single monophyletic *L. borgpetersenii* clade (Gomard *et al*. 2016; Dietrich *et al*. 2018). Interestingly, the screening of *M. acetabulosus* specimens through MAT reveals that sera agglutinate two distinct serogroups, *i*.*e*. Panama, and Pyrogenes. Although there is poor congruence between serogroups and genomospecies, members of Panama serogroup can be found in two species, *L. noguchii* and *L. inadai* (Levett 2001), but not in *L. borgpetersenii*. This suggests that the Panama serogroup signature results from independent systemic infections, which have cleared out without leading to renal colonization.

Although the molecular and serological analyses from *Rattus* and *Tenrec* were not all obtained from the same specimens, the results presented herein hardly support any agreement of data obtained from molecular and serological work. Interestingly, the investigation of these two mammal genera on Mayotte and Reunion Island highlight the importance of independently evaluating the reservoir status of a given mammal species on different islands, as *Rattus* do not shelter the same *Leptospira* species. The investigation of *Tenrec* is more compelling. While on Mayotte *T. ecaudatus* is the exclusive carrier of the recently described *L. mayottensis*, investigations on Reunion Island showed that up to 81% of individual *Tenrec* were seropositive (mostly reacting against Icterohaemorragiae) but not one individual showed evidence for chronic kidney infection (Desvars *et al*. 2013; Guernier *et al*. 2016). On Reunion Island, *T. ecaudatus* is therefore not a reservoir of *Leptospira* and Icterohaemorragiae serogroup revealed through MAT should be considered as evidence of animal exposure to the *Leptospira* actually present in the environment.

In our tested samples, some individuals were positive through PCR but negative through MAT. These discrepancies may result from past infections and subsequent kidney colonization followed by titer decay and eventually seronegativation, as previously reported in animal reservoirs (Lloyd-Smith *et al*. 2007) and incidental hosts (Blackmore, Schollum and Moriarty 1984; Lupidi *et al*. 1991; Cumberland *et al*. 2001). We propose that conflicting results of known reservoirs, such as bats testing positive through MAT but negative through PCR using kidney tissues and/or urine, are best explained by animals that experienced past infections in which *Leptospira* did not colonize the kidneys. This absence of colonization might be related to a low infecting dose and/or from an infecting bacterial genomospecies that is unable to establish persistent renal colonization in a specific vertebrate species. This assumption is based on the existence of host-*Leptospira* molecular determinants required for renal colonization, an hypothesis substantiated by experimental infections in which bat-borne and Tenrec-borne *Leptospira* were not able to lead to chronic infection in rats (Cordonin *et al*. 2020).

Finally, the biological setting of SWIO islands brings further evidence of problems using MAT for the identification of *Leptospira* reservoirs. Several studies have used MAT on samples of wild animals to address their role in the epidemiology of leptospirosis. As demonstrated here, this serological technique is very useful as it opens a window on environmental exposure to *Leptospira*. However, even though it is clearly important to address the diversity and intensity of *Leptospira* exposure in an environmental setting, we emphasize that MAT data from investigated animals cannot lead to any robust conclusion regarding their role as a reservoir. Such investigations, carried out in a One Health framework require bacterial genotyping using kidney or urine samples so that bacteria excreted by mammal reservoirs can be compared to those identified in acute human cases. Ultimately, a thorough investigation of leptospirosis following a One Health framework would require (i) PCR screening of urine or kidney tissues from putative animal reservoirs, (ii) isolation of *Leptospira* from identified reservoirs, and (iii) the inclusion of these isolates in a MAT panel used to screen human sera collected from persons living or having a professional/recreational activity in the investigated environment. Such investigations would allow not only identifying animal reservoirs in a specific environmental setting, but highlighting those bacterial species/lineages of major medical concern.

## Acknowledgements

The sampling of *Mormopterus acetabulosus* on Mauritius Island was supported by a grant from European Research Development Fund (ERDF) “Pathogènes associés à la Faune Sauvage Océan Indien” #31189. Sampling and export permits were granted by the National Parks and Conservation Service (NPCS) of the Ministry of Agro-industry & Food Security of Mauritius. Original serological data presented in the manuscript were granted by ERDF InterregV ECOSPIR “Eco-épidemiologie des leptospires endémiques de l’Océan Indien : des bactéries à risque pour les populations humaines ?” #RE6875. We are thankful to the director M. Puttoo and R. Sookhareea for their assistance in the field. This work is dedicated to the memory of Dr. A. Michault from the Centre Hospitalier Universitaire (CHU) Saint Pierre, who brought considerable help in the serological analysis.

---

**Table S1.** Details of *Leptospira* strains used for the Microscopic Agglutination Test on *Mormopterus acetabulosus*.

| Species | Serogroup | Serovar | Strain |
| --- | --- | --- | --- |
| *L. biflexa* | Semarang | Patoc | Patoc I (Paris) |
| *L. borgpetersenii* | Ballum | Castellonis | Castellon 3 |
|  | Sejroe | Hardjobovis | Sponselee |
|  | Sejroe | Sejroe | M 84 |
|  | Tarassovi | Tarassovi | Perepelicin |
| *L. interrogans* | Australis | Australis | Ballico |
|  | Autumnalis | Autumnalis | Akiyami A |
|  | Bataviae | Bataviae | Van Tienen |
|  | Canicola | Canicola | Hond Utrecht IV |
|  | Hebdomadis | Hebdomadis | Hebdomadis |
|  | Icterohaemorrhagiae | Copenhageni | Wijiberg |
|  | Icterohaemorrhagiae | Icterohaemorrhagiae | - |
|  | Pomona | Pomona | - |
|  | Pyrogenes | Pyrogenes | Salinem |
| *L. kirschneri* | Cynopteri | Cynopteri | 3522C |
|  | Grippotyphosa | Grippotyphosa | Moskva V |
|  | Mini | undertermined | 200803703 |
| *L. noguchii* | Panama | Panama | CZ 214K |

**Table S2.**
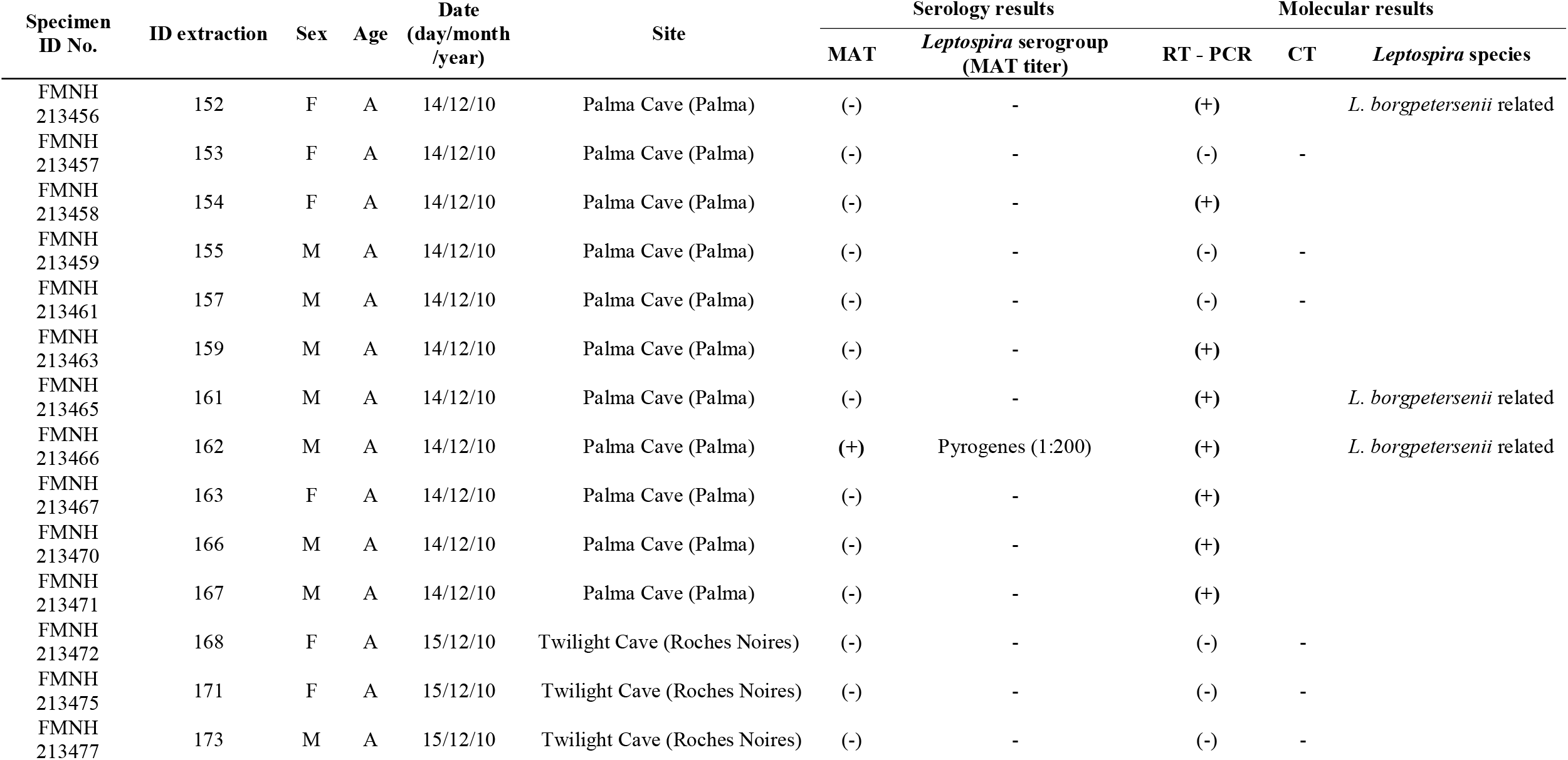
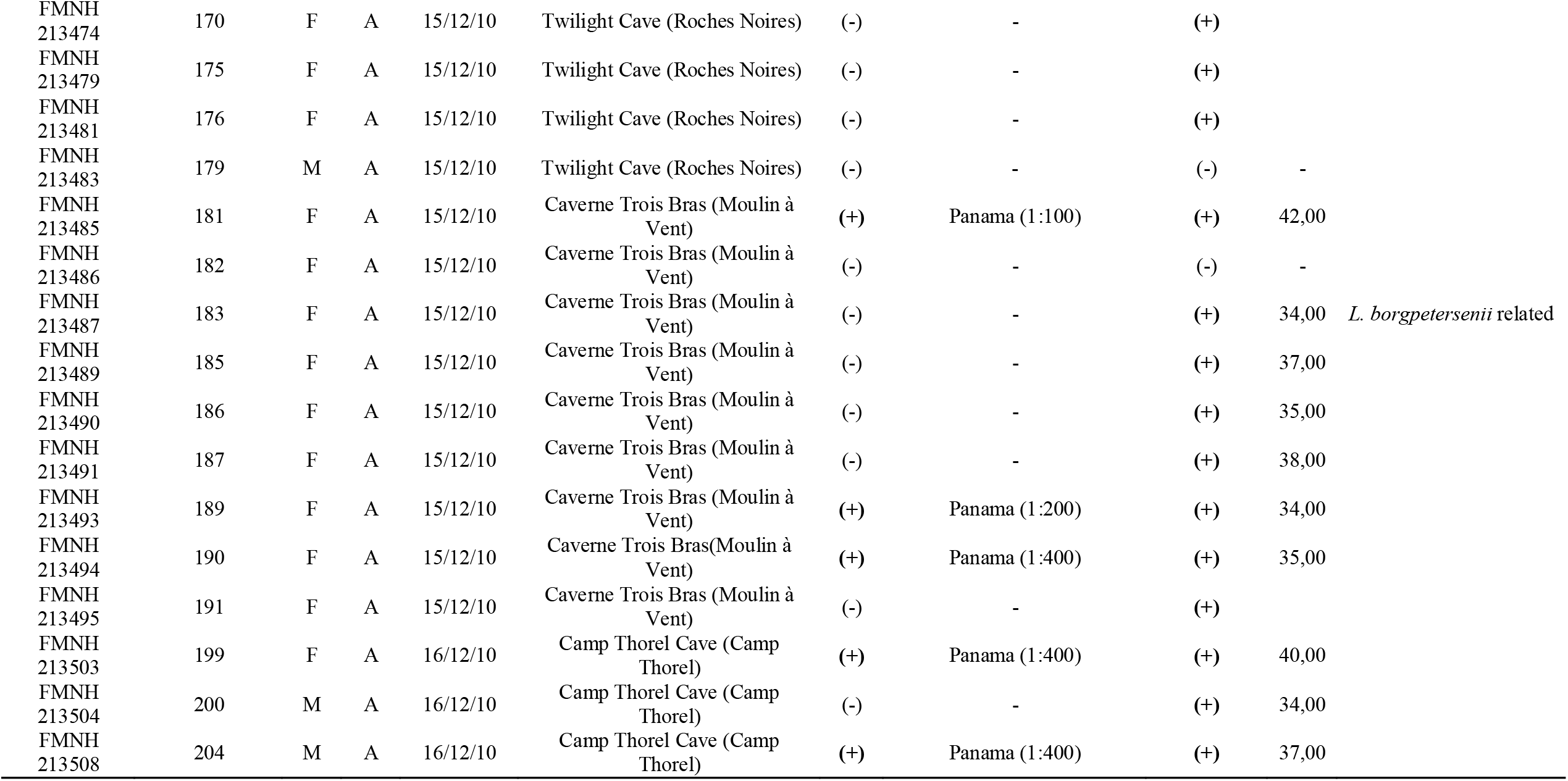
Details of bat samples, *Mormopterus acetabulosus* from Mauritius, used in the present study. The table includes the (a) Microscopic Agglutination Test (MAT) results and *Leptospira* molecular data obtained from the present study and the work of Dietrich et al. (2018) respectively. *RT-PCR: Real-Time Polymerase Chain Reaction, CT: Cycle Threshold, FMNH: specimen deposited in the Field Museum of Natural History*

## Notes

### Competing Interest Statement

The authors have declared no competing interest.

